# Hybrid breakdown is temporary and not expressed in a novel environment during 50 generations of experimental evolution in *Tetranychus urticae* hybrids

**DOI:** 10.64898/2026.07.16.738867

**Authors:** M. Kuijt, E. Villacís Pérez, S. Chakraborty, L. Dong, A. Wansink, S. Ebdon, K.S. Jaron, J. Kulmuni

**Affiliations:** Institute of Biodiversity and Ecosystem Dynamics, University of Amsterdam, Amsterdam, The Netherlands; Department of Ecology and Genetics, Uppsala University, Norbyvägen 18D, 75236 Uppsala, Sweden; Tree of Life, Wellcome Sanger Institute, Wellcome Genome Campus, Hinxton, Cambridge CB10 1SA, United Kingdom; Organismal and Evolutionary Biology, Helsinki University, Helsinki, Finland

## Abstract

Hybridization, the interbreeding between species or genetically distinct populations, can lead to deleterious fitness consequences, but simultaneously it can boost adaptive potential by increasing genetic variation, especially in novel environments. However, how incompatibilities and beneficial genetic combinations interplay across generations remains poorly understood. Here, we tracked how two fitness proxies, the absolute number of adult offspring and the proportion of eggs that reached adulthood, evolve in parental and hybrid populations at three different time points over 50 generations, in both novel and ancestral environments. To test this, we used two geographically distinct populations with low divergence (Dxy=0.002) of the two-spotted spider mite (*Tetranychus urticae*). Across the first three generations, hybrids showed significantly lower fitness than parental populations in the ancestral environment, indicating incompatibilities between the parental genomes. In contrast, hybrid and parental fitnesses were similar in the novel environment, indicating that the impact of incompatibilities was minor compared to the selection imposed by the novel environment. However, after 50 generations, hybrids displayed similar fitness relative to parental populations in all environments, suggesting resolution of the incompatibilities. Furthermore, around generation 45, hybrids temporarily outperformed parental populations in a novel environment, suggesting a transient window of higher adaptive potential, before fitness stabilized again by generation 50. In conclusion, we show that hybrid populations of *T. urticae* can swiftly purge incompatibilities when genetic divergence is low. These findings suggest that the dynamics of incompatibility resolution and adaptive potential of novel haplotypes play out over a long time frame, highlighting the importance of tracking hybrid fitness past the first few generations.

## Introduction

Hybridization, the interbreeding of two genetically distinct populations resulting in offspring of mixed ancestry (Barton & Hewitt, 1989), can lead to various consequences that can either reduce or increase individual fitness and population viability, eventually impacting the persistence of lineages over evolutionary timescales (Arnold, 1997; Seehausen, 2004; Abbott *et al*., 2013; Peñalba *et al*., 2024). A common consequence of hybridization is breakdown, where hybrids display lower fitness than parental populations due to genetic incompatibilities (Barton & Hewitt, 1985; Edmands, 1999; Willett, 2006; Stelkens *et al*., 2015; Thompson *et al*., 2024). These incompatibilities arise when alleles that evolved independently in allopatry interact negatively in hybrids, reducing offspring fitness, viability, or fertility (Dobzhansky, 1937; Muller, 1942). As genetic divergence increases, these incompatibilities are expected to accumulate, but even in early divergence incompatibilities may have large evolutionary consequences (Orr & Turelli, 2001; Ono *et al*., 2017; Coughlan & Matute, 2020). For example, mutations that reduce hybrid fitness can lead to selection favoring increased assortative mating (i.e. reinforcement) (Servedio & Noor, 2003; Butlin & Smadja, 2018).

However, hybridization can also lead to fitness benefits, and recent research has highlighted that hybridization is prevalent across taxa (Stelkens *et al*., 2015; Koevoets *et al*., 2012; Harris & Nielsen, 2016; Chaturvedi *et al*., 2020; Christe *et al.,* 2017). In the short term, hybridization can lead to heterosis, where first-generation offspring outperform one or both parental populations (Rieseberg *et al*., 1999). Over longer time scales, hybridization can facilitate faster adaptation due to the emergence of new genetic combinations (Meier *et al*., 2017), the increase in genetic variation (Satokangas *et al*., 2023), or the masking of recessive deleterious alleles (Crow, 1948; Whitlock *et al*., 2000). Eventually, hybrids can even evolve into entirely new species, boosting the generation of biodiversity (Rieseberg *et al*., 2003; Rosser *et al*., 2024).

Before the completion of speciation, hybridization between divergent lineages is likely to generate both deleterious and beneficial allelic combinations (Krapf *et al*., 2026). Over time, isolated hybrid populations are expected to purge incompatibilities, and while this depends on the recombination rate (Martin *et al*., 2019) as well as the severity and genomic architecture of incompatibilities, the rate of purging is not well understood (Blanckaert *et al*., 2021). Most empirical studies focus on the first few generations after hybridization (but see Mitchell *et al.,* 2019; Pinto & Stelkens, 2026), where incompatibilities are still segregating within large blocks of parental ancestry, resulting in varied outcomes, especially after the F1 generation (Edmands, 1999; Willett, 2006; Koevoets *et al*., 2012; Stelkens *et al*., 2015). However, theory and simulations predict that over generations, recombination breaks down these blocks (Buerkle & Rieseberg, 2008; Sachdeva & Barton, 2018) and hybrid fitness can recover and even exceed parental fitness (Kulmuni *et al*., 2023). Crucially, both theoretical and empirical studies suggest that hybrid populations especially may have an advantage over parental populations in novel environments where parents are not pre-adapted (Kulmuni *et al*., 2023; Thompson *et al*., 2024; Pinto & Stelkens, 2026). Using Fisher’s Geometric Model, Kulmuni *et al*. (2023) showed that simulated hybrid populations adapt faster than the parental populations in nearly all environmental scenarios, despite suffering from incompatibilities, except when the novel environment was close to the original environment of the parental populations. Here, we are interested in experimentally investigating the impact of incompatibilities over generations in the ancestral environment and the effect of hybridization on subsequent survival in novel environments.

The two-spotted spider mite (*Tetranychus urticae* Koch 1836) is a highly suitable model system to study proxies for fitness in hybrids over time and in different environments (Fry, 1989; Belliure *et al*., 2010). Because of their short generation time, they can be used to study the consequences of hybridization over tens of generations in real-time. Additionally, spider mites are haplodiploid, meaning that in haploid males, recessive incompatibilities are expressed and directly exposed to selection (Knegt *et al*., 2017; Nouhaud *et al*., 2020). Fitness proxies like egg-laying, hatching, and juvenile survival are easy to collect at the individual level, and experimental conditions can be replicated extensively due to their small size. Therefore, reproductive barriers can be tracked at a large scale (de Boer, 1982; Sato *et al*., 2015; Cruz *et al*., 2021). Moreover, investigating the adaptive potential of spider mites is crucial, as their ability to rapidly adapt to novel host plants or environments could exacerbate their impact as agricultural pests (Wybouw *et al*., 2015; Zhang *et al*., 2023). When different populations cross, hybridization can quickly spread adaptive traits across lineages (Sugasawa *et al*., 2002; Ito & Takatsuki, 2023; Xue *et al*., 2023; Villacis-Perez *et al*., 2025). Understanding these dynamics is therefore essential for predicting pest outbreaks.

Here, we established an experimental evolution experiment combined with whole genome sequencing of individual mites to answer the following questions: 1) Can two *Tetranychus urticae* populations, diverged in allopatry, hybridize and can the hybrids persist in lab conditions? 2) Is hybrid breakdown detected in hybrid populations of *Tetranychus urticae* and do the populations overcome hybrid breakdown during 50 generations? 3) Is there a difference between hybrid and parental populations in their ability to survive in novel environments and does this change over generations?

## Material and Methods

### Mite populations, rearing and plants

Two laboratory populations of *Tetranychus urticae* from geographically distinct origins were used to generate hybrid populations: ‘London’, originally collected from London, Canada (Grbic *et al*., 2011), and ‘Santpoort-2’ collected from Santpoort, the Netherlands (Kant *et al*., 2008). They have been maintained in the laboratory since 2011 and 2001, respectively. Both populations have since then been reared on detached leaves from common bean (*Phaseolus vulgaris* cv. Dubbele Witte zonder Draad) at 25°C, 60% relative humidity and a 16h:8h light:dark cycle.

We were interested in estimating fitness proxies in ancestral and novel environments. To do this, we used the ancestral host plant (common bean) and various novel host plants: quinoa (*Chenopodium quinoa* var. epidermal bladder cell-free (ebcf; Moog *et al*., 2022)), maize (*Zea mays*), and tomato (*Solanum lycopersicum,* var. Castlemart). Plants were grown in greenhouse conditions.

### DNA extraction and whole genome sequencing

To contextualize the genetic diversity of the focal populations (London and Santpoort - 2), individuals from two additional populations of *T. urticae* were sequenced: ‘Malaga’ (collected from Málaga, Spain in 2024) and ‘Red’ (collected from Miranda de Arga, Spain in 2018). *T. urticae* displays within-species color polymorphisms and has both green and red forms. Populations Santpoort-2, London, and Malaga belong to the green color morph and Red belongs to the red color morph. We sequenced genomes from 3 diploid female individuals from each population at Novogene (Germany). Additionally, three males and one female from the London population and two females from the Santpoort-2 population were sequenced at the Sanger Institute (Hinxton, UK).

To establish a cost-effective protocol for single-mite genomic analysis, we evaluated two DNA extraction methods: ‘SOP – Lysis C plate-based DNA extraction’ (Korlevíc *et al*., 2023) and the PicoPure DNA Isolation Kit (ThermoFisher). Both methods yielded comparable DNA concentrations (Supplementary Figure 5). Therefore, the more cost-effective method, i.e. Lysis C, was chosen for all downstream extractions. Briefly, this method uses an in-house lysis buffer (for 100 μl buffer: ddH_2_O 72.95 μl, Tris 20 μl, EDTA 5 μl, Tween-20 0.05 μl and Proteinase K 2 μl). Single spider mite individuals were crushed in 20 μl lysis buffer and subsequently incubated at 55℃ overnight. After incubation, samples were transferred into new tubes. Commercial provider Novogene (Germany) performed library preparation and subsequent whole-genome sequencing. Briefly, DNA was fragmented, end-polished, A-tailed, and ligated with Illumina adapters. Sequencing was performed on an Illumina NovaSeq X Plus Series, generating 150 bp paired-end reads and resulting in 17-25 million reads per sample. The sequencing at the Sanger Institute was also performed on an Illumina NovaSeq X Plus Series, generating 150 bp paired-end reads and resulting in 12-73 million reads per sample.

### Sequence filtering, analysis, and statistics

Raw sequencing reads and adapter sequences were trimmed and filtered using Trimmomatic v0.39 (parameters LEADING: 3, TRAILING: 3, MINLEN: 100). Subsequently, the reads were aligned to the *T. urticae* reference genome from Cao *et al*. (2024; Accession: GCA_036877765.1) using bwa mem (version 0.7.17), using default parameters. Duplicates were removed using Picard’s MarkDuplicates (version 2.26.1). Unmapped reads and reads with a quality score below 20 were filtered using samtools (version 1.23.1). Bcftools mpileup (version 1.17) was used for single nucleotide polymorphism (SNP) calling and lastly, SNPs were filtered using bcftools, keeping only biallelic SNPs with a quality score of 30 or higher, a maximum of 5% missing data across samples, and a minor allele frequency equal to or higher than 5%. This pipeline resulted in a total of 1.1 million high-quality, genome-wide SNPs.

Nucleotide diversity and divergence between the parental populations were calculated using sliding window-based population statistics using pixy, specifically nucleotide diversity (π), absolute sequence divergence (Dxy), and the fixation index (Fst), using 5 kb windows with a 2.5 kb step. Additionally, a Principal Component Analysis (PCA) was conducted in R (Version 4.5.0).

### Generation of hybrid populations

Hybrid populations were made following Godinho *et al*. (2020), resulting in four experimental populations: two parental populations (London and Santpoort-2) and two hybrid populations, which resulted from crossing London females with Santpoort-2 males (LS) and vice versa (SL). Bidirectional crossing was done to take into account the effects of potential cytoplasmic incompatibility (CI) induced by *Wolbachia* (Vala *et al*., 2000).

First, 60 females from each parental population were placed on detached leaves of common bean to oviposit for 72H. The egg cohorts were used to generate individuals of a similar age. From the resulting offspring, 60 virgin females were isolated and crossed with 60 males from the alternative population, on individual leaf discs. Pairs were allowed to mate for 72H, after which the parental mites were removed, and the offspring could develop further. Since spider mites are haplodiploid and males are born from unfertilized eggs from the mother, only the females from this initial crossing were hybrids. Therefore, males were removed before the F1 females could mate with them, resulting in virgin F1 hybrid females. Consequently, these F1 hybrid females produce only hybrid males. Sixty F1 hybrid females were crossed with hybrid males, which were not their sons, to produce hybrids with on average 50:50 ancestry from both parental populations.

### Experimental setup

Parental and hybrid populations were initially maintained on both common bean and quinoa for the first ∼14 generations, in three replicate populations. Due to limited resources in rearing for colony maintenance on quinoa, replicates were combined and both parental and hybrid populations were maintained exclusively on bean after the 14th generation. To estimate fitness proxies over generations, fitness assays were performed across three different time points after hybridization: the first three generations (hereafter ‘early generation hybrids’ (EGH)), around generation ∼45 (‘late generation hybrids I’ (LGH I)) and around generation ∼50 (‘late generation hybrids II’ (LGH II)). Fitness proxies were measured either on the ancestral host (common bean) or a novel host (quinoa, maize, or tomato, depending on the experiment). Between the EGH experiments that were performed in 2023 (generations F1-3) and the LGH experiments performed in 2024/2025 (F45, F50), all populations were reared on detached bean leaves at 25°C, 60% relative humidity and a 16h:8h light:dark cycle.

We were interested in how the fitness of hybrids in relation to parents changed over generations in novel environments. In the EGH, fitness proxies for hybrids and parental populations were therefore evaluated on three-week-old quinoa as a novel host. In the LGH I, fitness proxies were evaluated on two-week-old maize due to the temporary unavailability of quinoa. In the LGH II, fitness proxies were consecutively evaluated on quinoa, maize, and tomato. All experiments across all generations were performed alongside the ancestral host (common bean).

**Table 1:** Overview of different experiments performed in this research.

| Evolutionary Window | Generations | Populations were grown on | Populations Tested | Host plants on which fitness proxies were evaluated |
| --- | --- | --- | --- | --- |
| Early-Generation Hybrids (EGH) | F1-F3 | Bean and quinoa | London, Santpoort-2, 2 Hybrid populations | Common Bean (Ancestral)<br>Quinoa (Novel) |
| Late-Generation Hybrids I (LGH I) | F45 | Bean | London, Santpoort-2, 2 Hybrid populations | Common Bean (Ancestral)<br>Maize (Novel) |
| Late-Generation Hybrids II (LGH II) | F50 | Bean | London, Santpoort-2, 2 Hybrid populations | Common Bean (Ancestral)<br>Quinoa (Novel)<br>Maize (Novel)<br>Tomato (Novel) |

### Fitness proxies

To estimate individual fitness of the two-spotted spider mites, the following fitness proxies were used: 1) the absolute number of adults, referring to the number of adults that resulted from the offspring of a single mated female by day 12, which was chosen to indicate population growth, and 2) the proportion of eggs that reached adulthood, referring to the proportion of eggs that had developed into adults by day 12 per single mated female. This proxy was chosen to estimate the developmental success of the offspring.

Fitness proxies were measured using individual mite pairings on leaf discs of 1.5 cm, which were maintained on wet cotton. For the fitness proxies, a single virgin female of the same age and one adult male were collected from the experimental populations and placed on individual leaf discs (EGH: n = 20 per replicate, experimental population, host plant, and generation, EGH total n = 1440, LGH: n = 20 per experimental population and host plant, LGH I total n = 160 and LGH II total n = 480). Collectively, fitness proxies were measured for 2080 mite pairings over the entire experiment.

#### Early Generation Hybrids

To measure the fitness proxies in the EGH, parents were removed from leaf discs after a 72H period, and the number of eggs laid per female and parental survival was recorded. Over the subsequent 12 days, offspring development and survival were measured on days 1, 3, 5, 8, 10 and 12.

#### Late Generation Hybrids I (LGH I)

During the LGH I experiment (generation ∼45), fitness proxies were recorded following the same experimental procedures as described for the EGH.

#### Late Generation Hybrids II (LGH II)

Fitness proxies were recorded with a modified protocol for the LGH II experiment (generation ∼50). Mites were first allowed to acclimatize to the new host plant for a 48-hour period before the mating and egg-laying timeline began, in contrast to both the EGH and LGH I experiments. This step was added to reduce initial shock effects on the novel host and to minimize maternal environmental effects. All eggs laid during this initial 48-hour window were removed from the leaf discs. Following this acclimatization phase, the protocol continued as described before: after a 72H oviposition period, the parents were removed and the total number of eggs was counted. Further fitness measurements were performed as in the EGH.

### Statistical analysis

For the first fitness proxy, the absolute number of adults that emerged from the offspring, data were analyzed using a GLM with a Poisson distribution. In cases of significant overdispersion, a Quasipoisson distribution was used. The second fitness proxy, the proportion of eggs that reached adulthood, was analyzed using a GLM with a quasibinomial distribution, again to account for overdispersion. Post-hoc pairwise comparisons and odds ratios were calculated in R using emmeans from the ‘emmeans’ package (Searle *et al*., 1979; Lenth *et al*., 2026).

Because of slight methodological adjustments in the estimation of fitness proxies in EGH, LGH I and LGH II (e.g. acclimatization phase and different seed batches of host plants), absolute values of the fitness proxies were not compared across these generations.

## Results

### WGS from single mites yields reliable data

Of every individual mite, between 20 and 50 ng total DNA was extracted, with a genomic DNA fragment size peak over 15000 bp (Supplementary Figure 4), indicating high molecular weight DNA. Whole Genome Sequencing across 15 female samples from four populations was performed, resulting in an average coverage of 36x per individual. Additionally, 3 males were sequenced, with an average coverage of 96x per individual.

### London and Santpoort-2 populations are genetically distinct, although closely related

The genetic relationships among four *T. urticae* populations were investigated using PCA, based on 1.1 million genome-wide SNPs (Figure 1a+b). PC1 and PC2 accounted for 50.84% and 28.95% of the total variance, respectively. The London and Santpoort-2 populations form a distinct cluster relative to both the Red and Malaga populations. PC3 accounted for 6.66% of the variation, separating London and Santpoort-2. Pairwise genetic distance analyses revealed differentiation and divergence throughout the genome between the London and Santpoort-2 populations (Fst: 0.30, Dxy: 0.002), albeit still much lower than the differentiation and divergence between Santpoort-2 - Malaga (Fst: 0.82, Dxy: 0.011) and Santpoort-2 - Red (Fst: 0.75, Dxy: 0.008).

**Figure 1:**
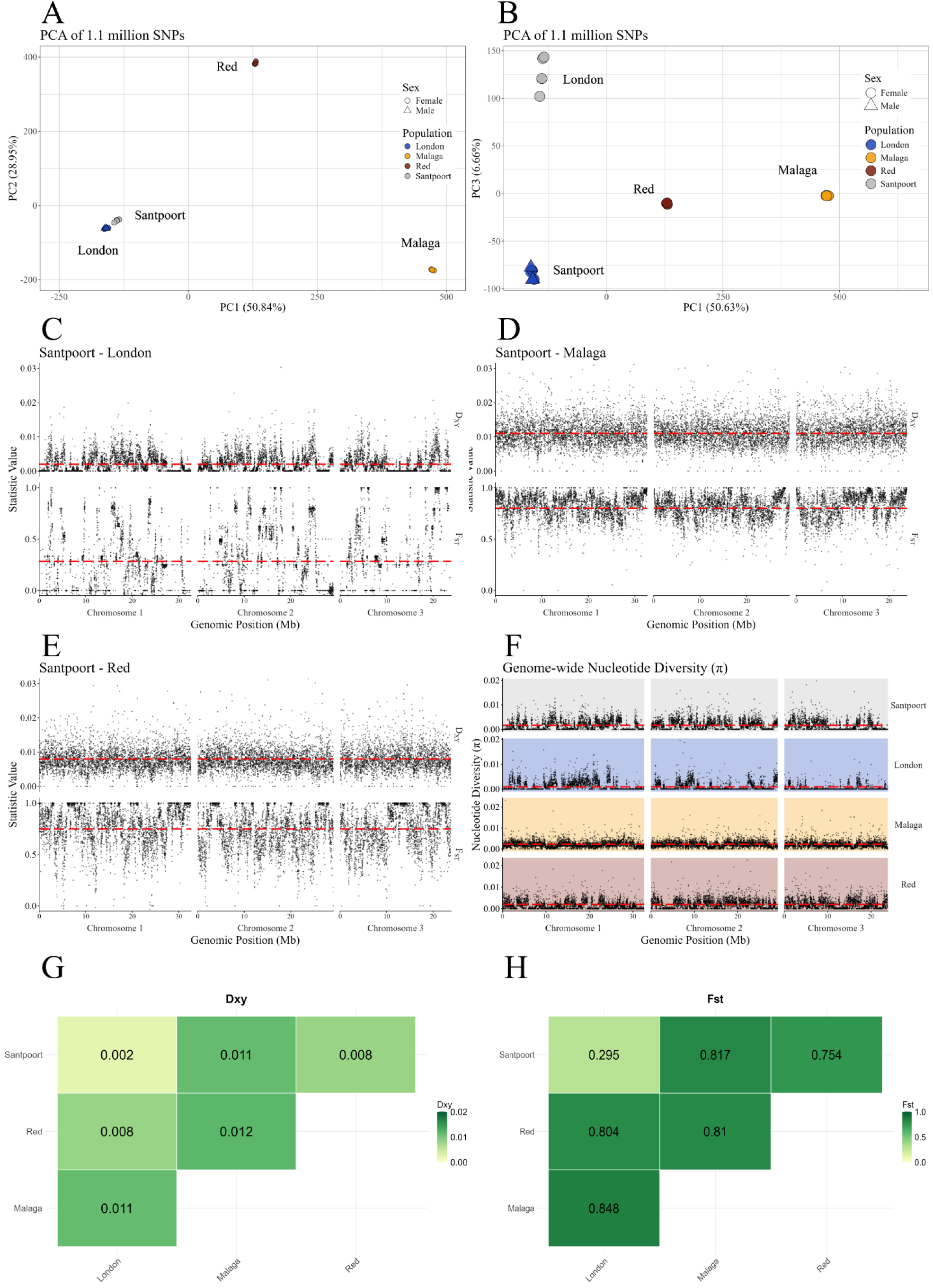
Extent of genetic differentiation, divergence and nucleotide diversity among four *Tetranychus urticae* populations. (a-b) Genetic differentiation among all populations (n = 17) as assessed by a Principal Component Analysis (PCA) of 1.1 million genome-wide SNPs. Different colours reflect different populations. (c-e) Genome-wide divergence (Dxy) and differentiation (Fst) between Santpoort-2 and three different spider mite populations: c) Malaga, d) London, and e) Red. In each panel, top graphs show absolute divergence (Dxy) values and bottom graphs show fixation index (Fst) values across the genome in 5 kb sliding windows based on 1.1 million SNPs (n = 3 diploid genomes per population). The red dashed lines show the genome-wide averages, as also summarized in the Fst and Dxy heatmaps in (g-h). Darker colours indicate higher values. (f) Genome-wide nucleotide diversity (π) values of all four populations.

To characterize the within-population diversity of all four populations, genome-wide nucleotide diversity (π) was calculated. While the Malaga and Red populations show high and relatively uniform diversity across all three chromosomes (average π = 0.13 and π = 0.10, respectively), London and Santpoort-2 show lower and more heterogeneous diversity (average π = 0.04 and π = 0.08, respectively).

Notably, when looking at between-population differentiation across the genome (Figure 1c-h), Santpoort-2 - London shows a highly variable differentiation (Figure 1c). In contrast, both Santpoort-2 - Malaga (Figure 1d) and Santpoort-2 - Red (Figure 1e) comparisons show more even differentiation across all chromosomes, especially when looking at the Dxy values, indicating strong divergence between the populations.

### Hybrid breakdown is detected in early generation hybrids, but it is not expressed on a novel host plant

Hybrid populations showed significantly reduced fitness relative to parental populations on the ancestral host (bean) in the first three generations after hybridization (i.e. EGH). This is seen as a 29% reduction in the absolute number of adults (mean difference in the absolute number of adults = 7.1 ± 2.12 SE, p < 0.01, GLM) and a 33% reduction in the proportion of eggs that reached adulthood (parental-to-hybrid odds ratio of proportion of eggs that reached adulthood = 2.9 ± 0.417 SE, p < 0.001, GLM). The reduction in fitness in hybrids compared to parental populations confirms that despite low overall divergence of the two populations, incompatibilities segregate in the hybrid genomes. There was no consistent significant difference in fitness proxies between the different hybrid populations (SL and LS), suggesting maternal effects like *Wolbachia* do not play a role in hybrid incompatibilities between the populations used (Supplementary Figures 1-3).

Interestingly, there was no difference between parental and hybrid populations in fitness proxies on the novel host quinoa (mean difference in the absolute number of adults = 1.47 ± 1.56 SE, p = 0.78; parental-to-hybrid odds ratio of the proportion of eggs that reached adulthood = 0.95 ± 0.20, p = 0.99). However, the fitness of parents was significantly lower in novel host quinoa compared to the ancestral host (common bean) (mean difference in the absolute number of adults = 14.34 ± 1.88 SE, p < 0.001; ancestral-to-novel host odds ratio of the proportion of eggs that reached adulthood = 3.70 ± 0.62, p < 0.001). This suggests that the impact of hybrid incompatibilities is relatively small compared to the strong fitness reduction caused by the novel environment.

**Figure 2:**
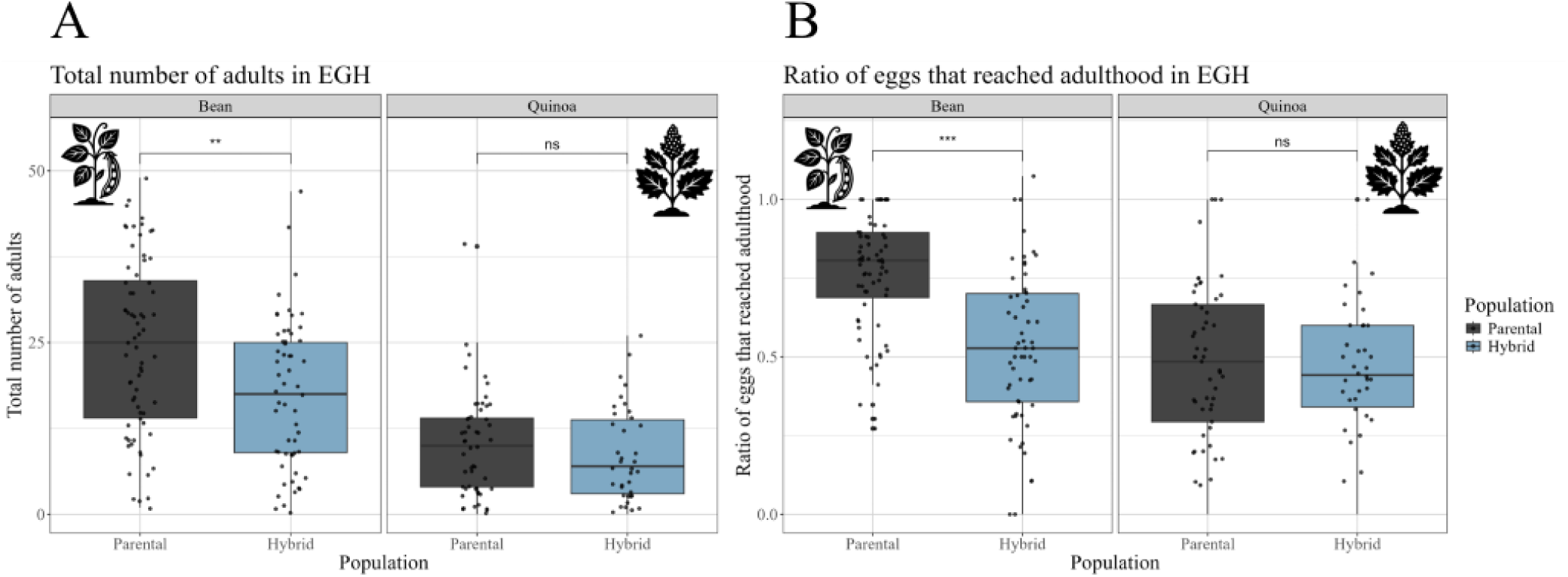
Fitness measures of parental populations (London and Santpoort-2) and early generation hybrids (EGH, F1-F3), tested on the ancestral (bean) and a novel host plant (quinoa). a) The absolute number of adults; b) The proportion of eggs that reached adulthood (n = 40 per host plant, per population, and generation).

### Hybrids recover from breakdown in later generations

After a period of 1.5 years, during which all populations were maintained on the ancestral host (bean), the fitness of hybrid and parental populations was reassessed. The proportion of eggs that reached adulthood and the absolute number of adults were no longer significantly different between hybrid and parental populations on the ancestral host bean in LGH II (generation ∼50) (Figure 3; Supplementary Figure 3), suggesting that hybrid incompatibilities are resolved.

**Figure 3.**
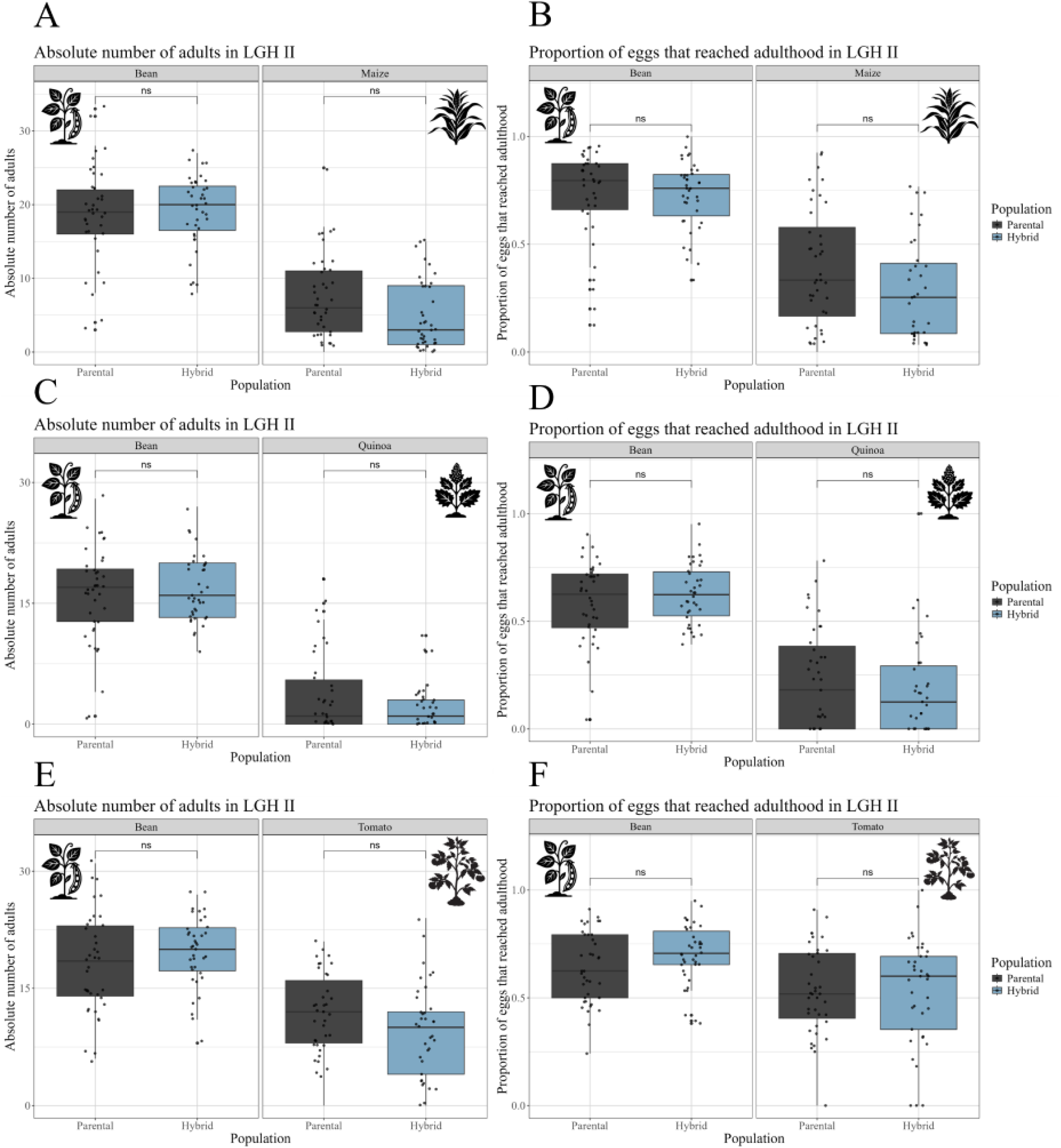
Fitness measures of Late Generation Hybrids II (LGH II) parental populations (London and Santpoort-2) and hybrids, tested on the ancestral (bean) and three different novel host plants: (a + b) maize, (c + d) quinoa and (e + f) tomato. Left panels (a, c, and e) show the absolute number of adults. Right panels (b, d, and f) show the proportion of eggs that reached adulthood (n = 40 per host plant, per population).

### Minor adaptive potential in Late Generation Hybrids

Interestingly, around generation 45 (LGH I), where hybrids and parents were tested on maize as a novel host, hybrids produced 42.6% more adults compared to the parental populations (parental-to-hybrid odds ratio of proportion of eggs that reached adulthood = 1.43 ± 0.17 SE, p-value < 0.05, GLM) (Figure 4a). At around 50 generations (LGH II), this adaptive potential of hybrids seems to have disappeared again as the fitness of hybrid and parental populations is comparable on three different novel hosts (maize, quinoa, tomato; Figure 3). This transient window of adaptive potential is in line with previous simulations (Kulmuni *et al*., 2023), where it was shown that hybrid fitness increases faster over generations than parental fitness in a novel environment, but only until a certain time frame after the hybridization event. Note, our experiment differs from the simulations in that we did not keep hybrids in the novel environment for the entire duration of the experiment.

**Figure 4:**
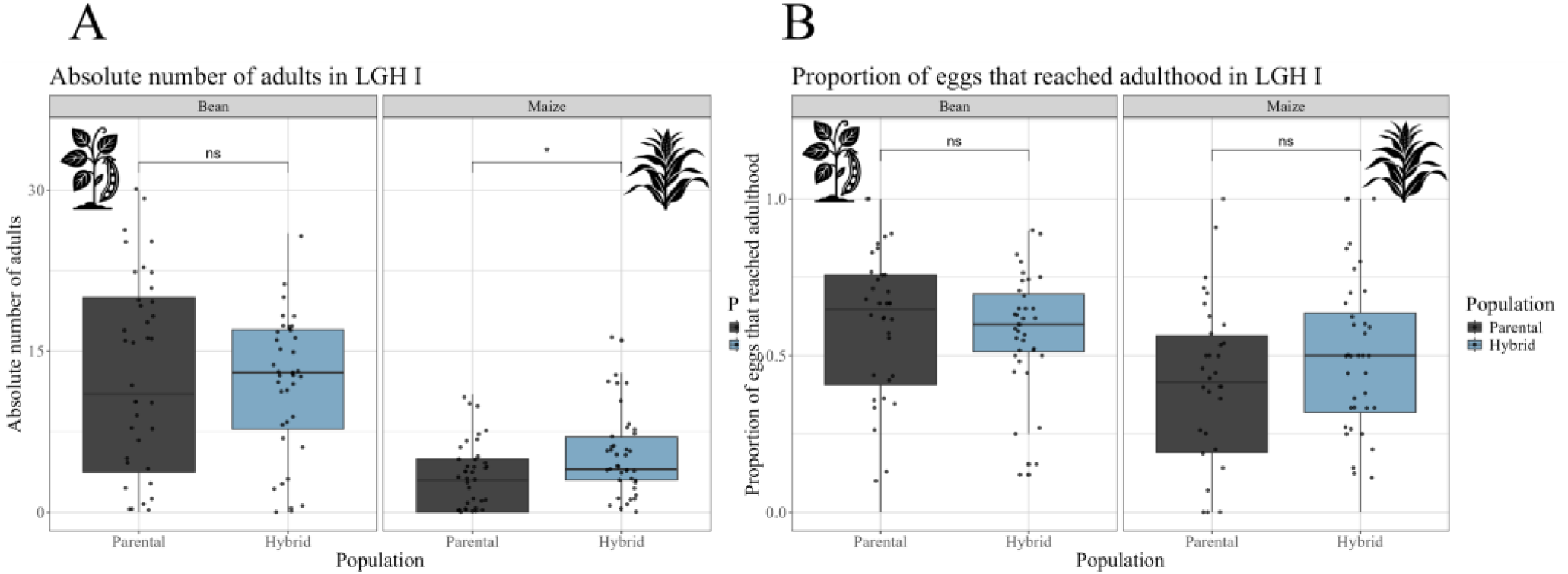
Fitness measures of Late Generation Hybrids I (LGH I, generation ∼45) parental populations (London and Santpoort-2), tested on the ancestral (bean) and a novel host plant (maize). a) The absolute number of adults b) The proportion of eggs that reached adulthood (n = 40 per treatment, per population).

## Discussion

Hybridization is an evolutionary driver that can cause breakdown due to genomic incompatibilities, but how these behave over time and across different environments remains poorly understood (Kulmuni *et al*., 2020; Blanckaert *et al*., 2021; Thompson *et al.,* 2024; Krapf *et al.,* 2026). Here, we show that genomic incompatibilities expressed in the first three generations are purged over 50 generations of experimental evolution when divergence between hybridizing populations is low. Furthermore, the impact of incompatibilities is minor compared to the maladaptation caused by a novel environment. These results align with findings from our previous simulations (Kulmuni *et al*., 2023) and have important implications for the outcomes of hybridization in a changing environment.

Firstly, we demonstrated that two genetically divergent populations of *T. urticae*, London and Santpoort-2 with Fst = 0.3 and Dxy = 0.002, can successfully hybridize and persist in laboratory conditions, despite an initial 29% reduction in fitness. The overall degree of divergence between the hybridized populations is comparable to within-species divergence that results in sterility or inviability in other taxa (Martin *et al*., 2013; Mérot *et al*., 2017; de Jong *et al*., 2025), but similar divergence values are also observed between young species with genome-wide barrier effects, like the mound-building wood ants (Satokangas *et al*., 2026; Heidbreder *et al*., 2026). The extent of early-generation hybrid breakdown observed in this study aligns with other systems, like cichlid fish (Stelkens *et al.,* 2015) and parasitic wasps (Koevoets *et al.,* 2012). However, this reduction in fitness seems to be transient: hybrid fitness recovered to parental fitness by generation 50, indicating purging of incompatibilities. In haplodiploid organisms like *T. urticae,* purging might happen even faster than in diploid organisms, since recessive allele combinations are directly exposed to selection in haploid males and cannot be masked (Nouhaud *et al*., 2020).

Between the London and Santpoort-2 populations, genetic differentiation is highly heterogeneous throughout the genome, with some genomic regions showing very low divergence. The underlying causes for the regions of low divergence are unknown, but they could be a result of low recombination, past gene flow between the populations or lab adaptation, since both populations have been kept in laboratory conditions for many years. Further studies are needed to elucidate the underlying factors of heterogeneous divergence and the potential role of recombination and selection.

When considering parental and hybrid fitness in a novel environment, the hybrid breakdown observed in early generation hybrids is masked by the pressures from this novel environment, resulting in equal fitness for parental and hybrid populations. This pattern supports the notion that ecological conditions can modulate the expression of genetic incompatibilities (Kulmuni & Westram, 2017; Thompson *et al*., 2022, 2024; Kulmuni *et al.,* 2020). Interestingly, by generation ∼45, hybrid populations seem to exceed parental fitness temporarily in the novel environment. However, this indication of hybrid vigor has disappeared by generation ∼50, suggesting that while hybrids may display some adaptive potential by generating new allelic combinations (Rieseberg *et al.,* 2003; Meier *et al*., 2017; Mitchell *et al.,* 2019), this genetic variation eventually subsides. Likely, this is because fixation occurs in the beneficial loci, in turn reducing adaptive potential (Kulmuni *et al*., 2023), even though we did not keep hybrids in the novel environment for the entire duration of the experiment (i.e. between generations ∼14 and ∼45).

Our findings have important implications for natural populations. Global change is expected to facilitate hybridization by bringing previously isolated populations into contact while simultaneously subjecting these populations to novel environmental conditions through climatic change and anthropogenic factors. Our results suggest that under these conditions, the impact of hybrid incompatibilities may be smaller than previously estimated. Importantly, hybrid populations can also overcome these incompatibilities, although this process will depend on the severity and complexity of the incompatibilities. Future experiments need to investigate hybrid fitness in different environments over the time frame between the 4th and 40th generations to further understand the interplay between incompatibilities and adaptive potential in hybrid populations.

## Acknowledgements

We thank colleagues at all Plants ‘n Bugs and SpeciAnt meetings for helpful discussions and feedback.

