## Supplementary Figure for "Hybrid breakdown is temporary and not expressed in a novel environment during 50 generations of experimental evolution in *Tetranychus urticae* hybrids"

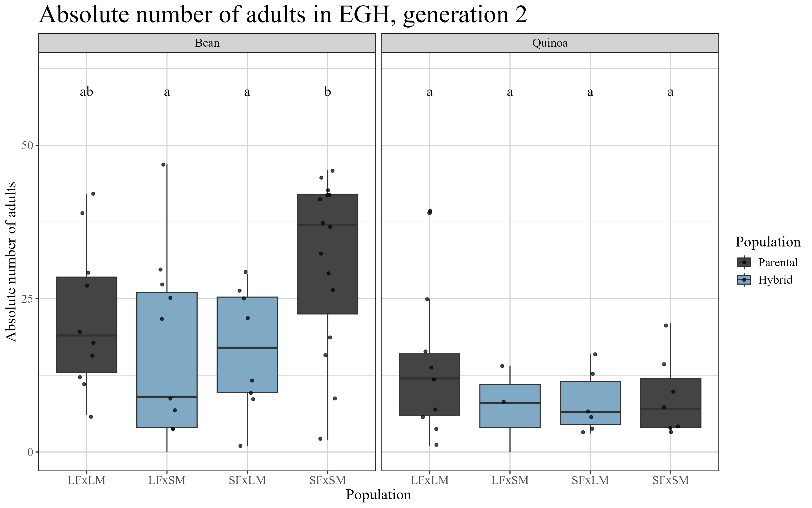

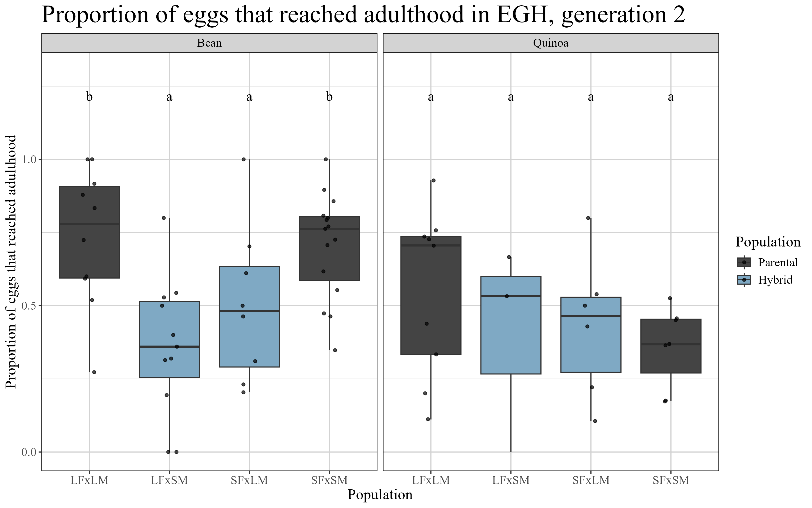

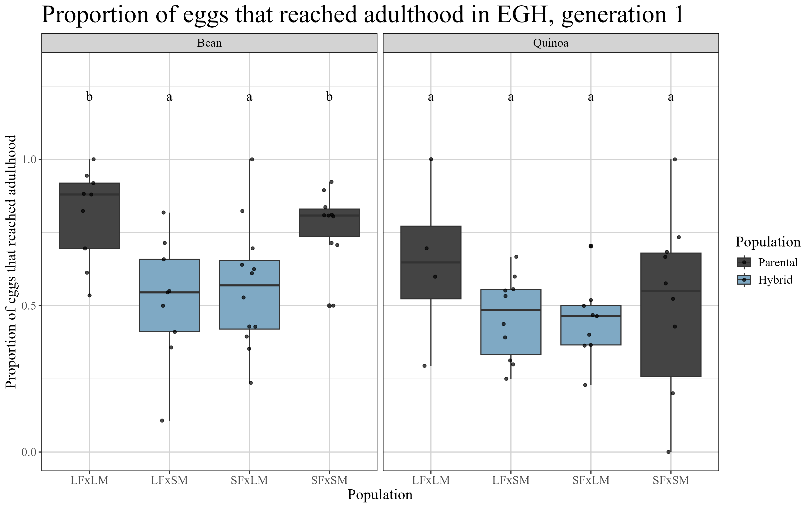

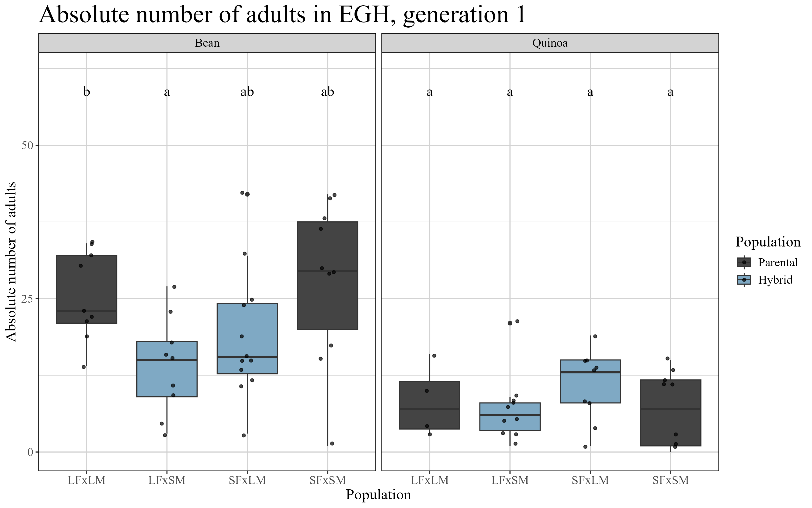

C

D

A

B

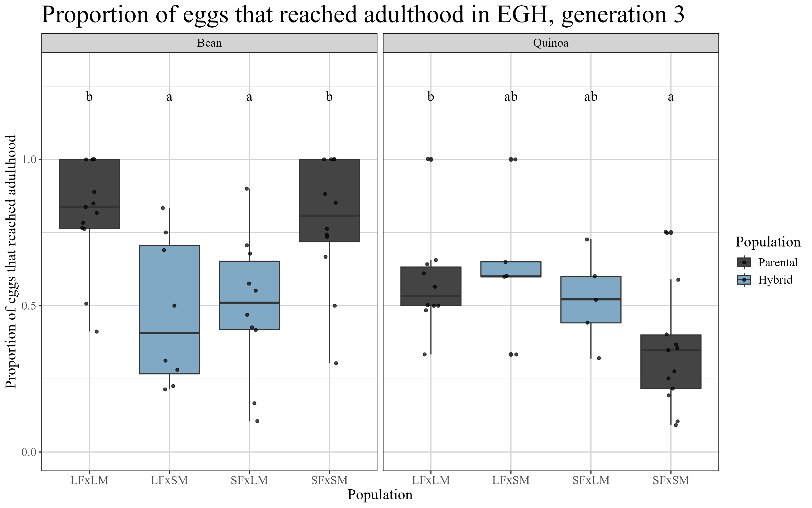

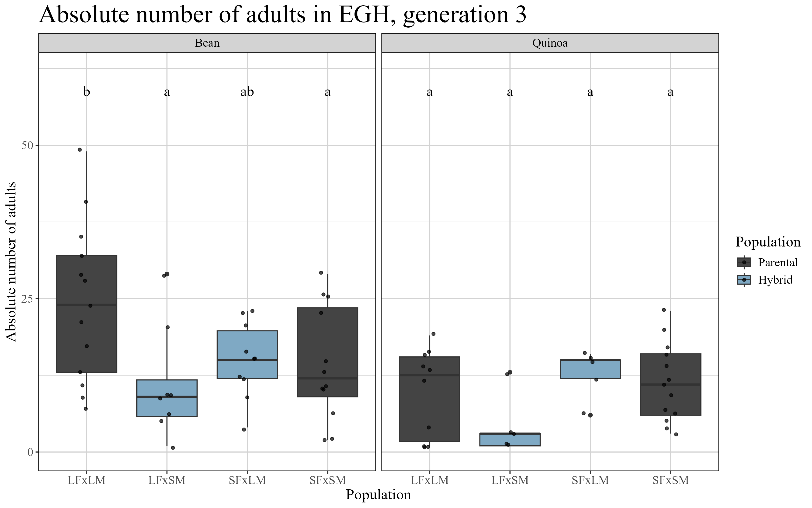

E

F

Supplementary Figure 1: Fitness measures of parental lines (LFxLM and SFxSM) and Early Generation Hybrids (EGH) visualised for both crossing directions (F1-F3, LFxSM and SFxLM), tested on the ancestral (bean) and a novel host plant (quinoa). Measures are averages over first 3 generations.

a + b) generation 1, c + d) generation 2 and e + f) generation 3. Left panels (a, c, and e) show the absolute number of emerged adults. Right panels (b, d, and f) show the ratio of eggs that reached adulthood, calculated as the number of adults divided by the number of eggs initially laid (n = 20 per treatment). Different letters indicate statistically significant differences (p < 0.05).

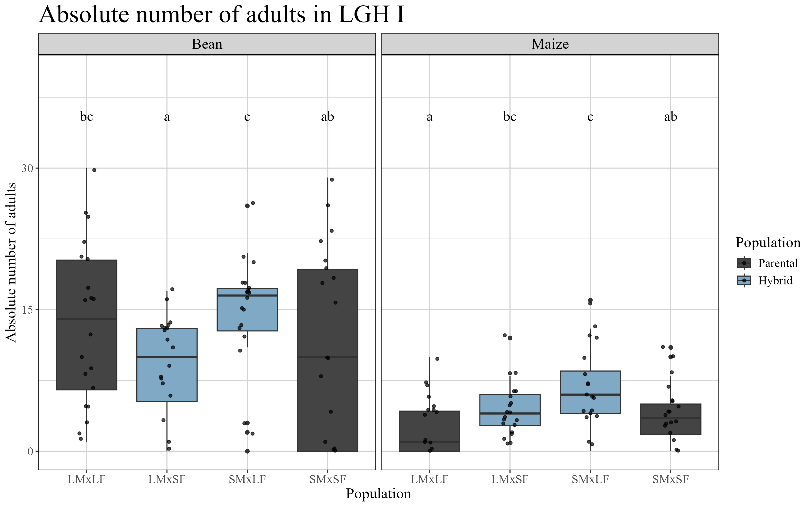

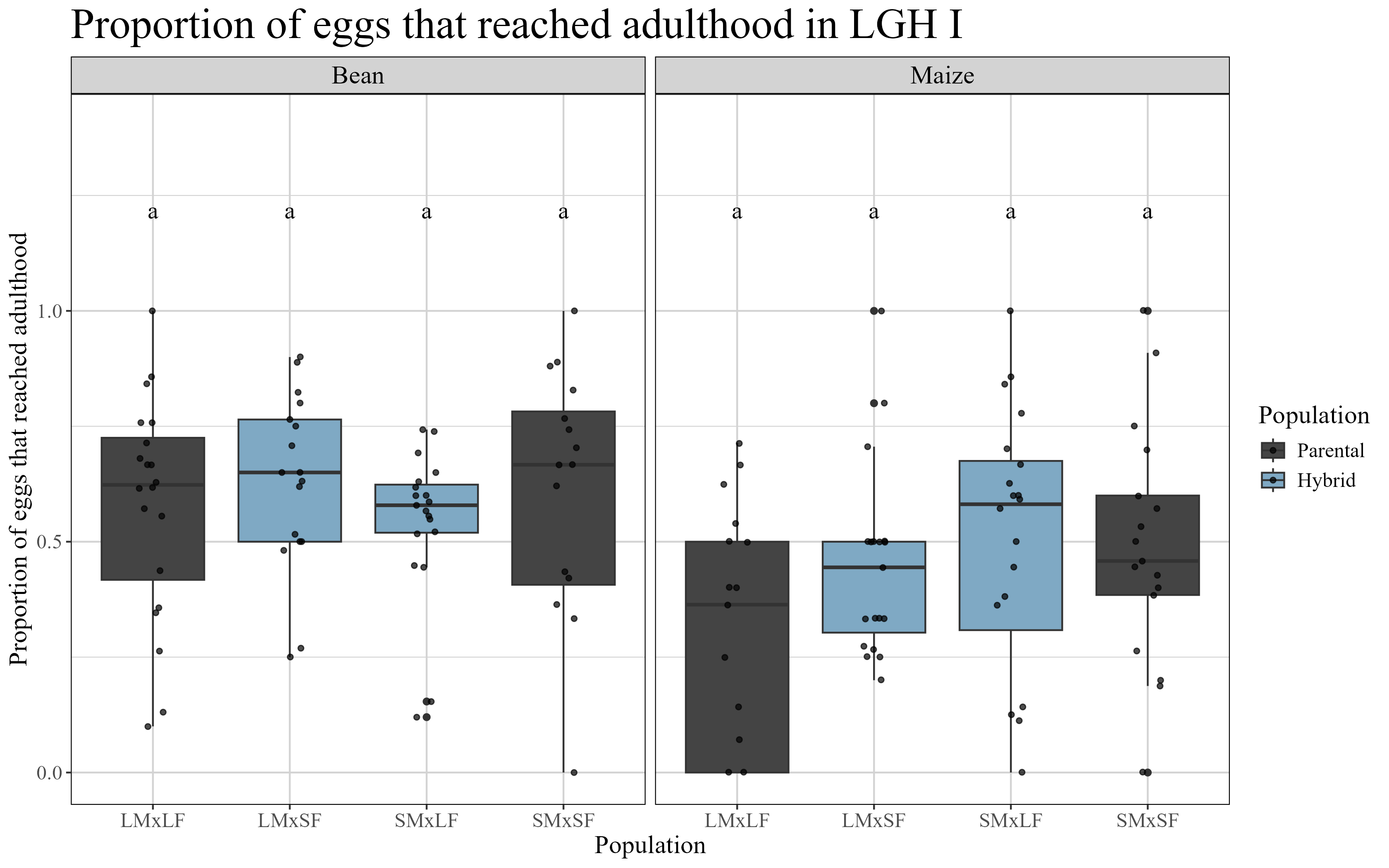

A

B

Supplementary Figure 2: Fitness measures of parental lines (LMxLF and SMxSF) and Late Generation Hybrids (LGH I, generation 45) visualised for both crossing directions (LMxSF and SMxLF) tested on the ancestral (bean) and a novel host plant (Maize). a) The absolute number of emerged adults b) The ratio of eggs that reached adulthood, calculated as the number of adults divided by the number of eggs initially laid (n = 20 per treatment). Different letters indicate statistically significant differences (p < 0.05).

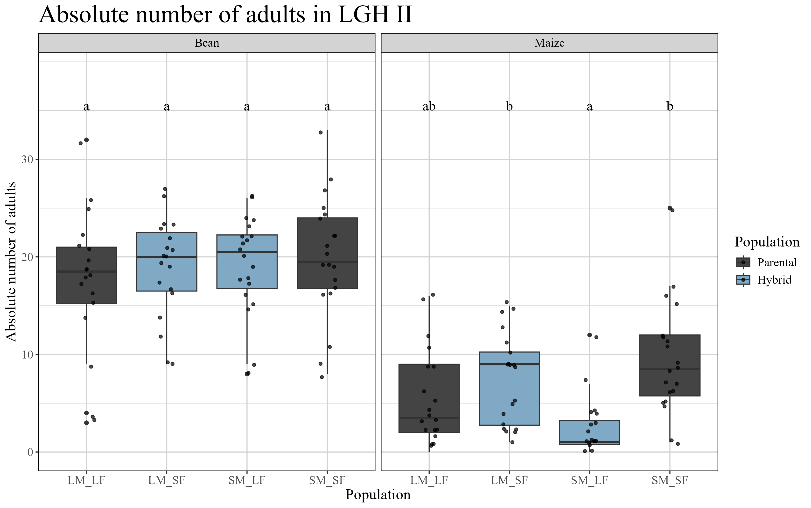

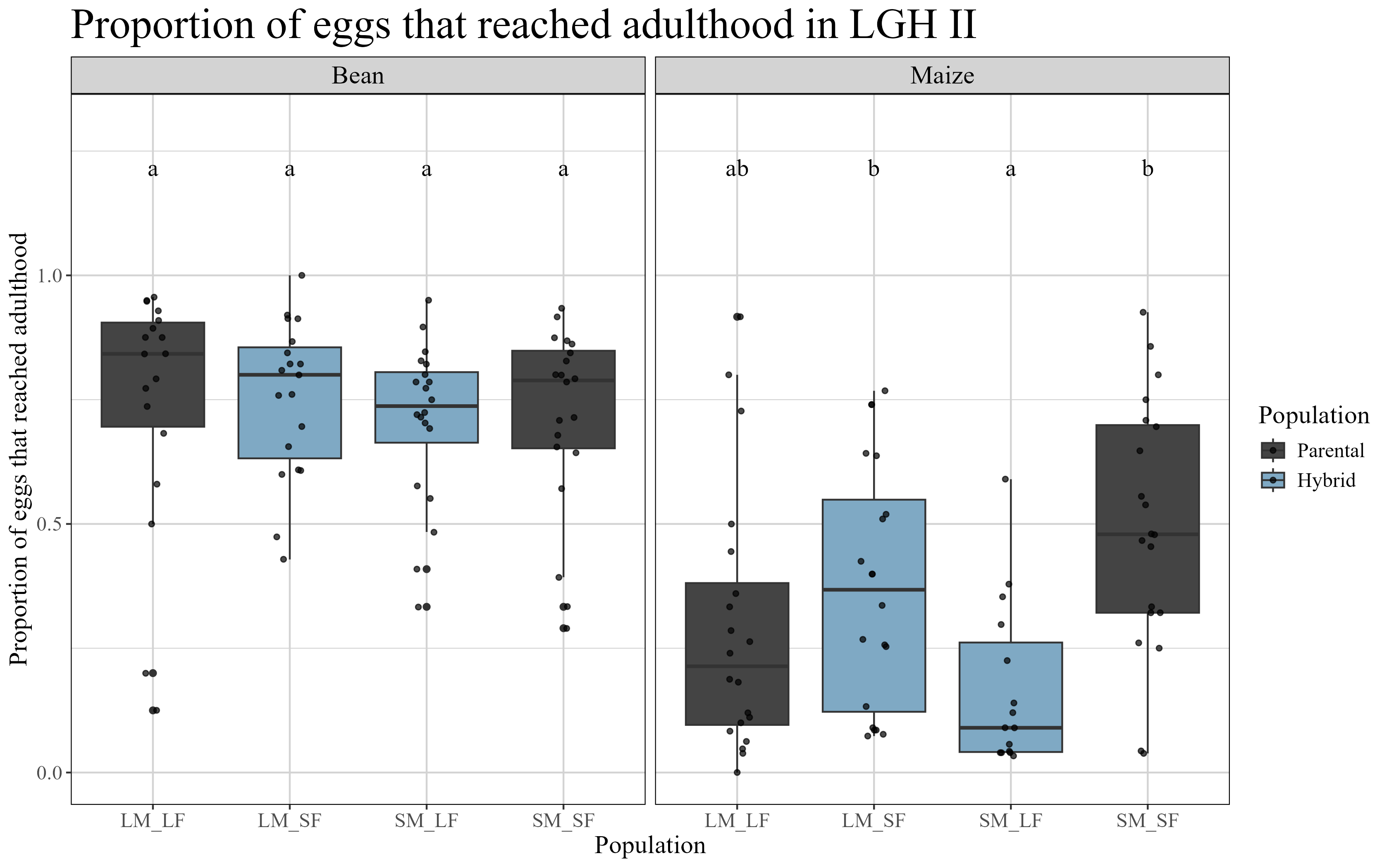

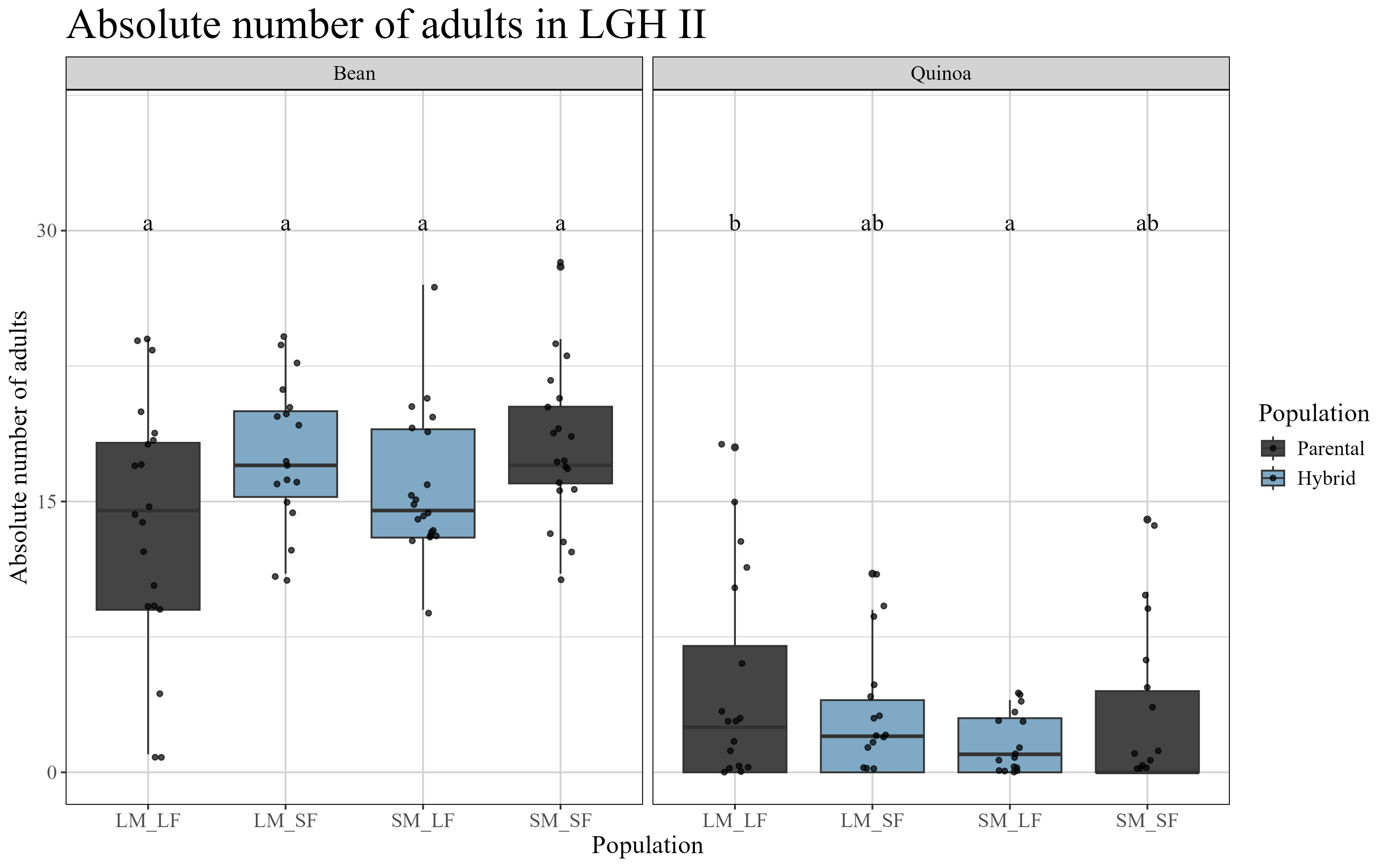

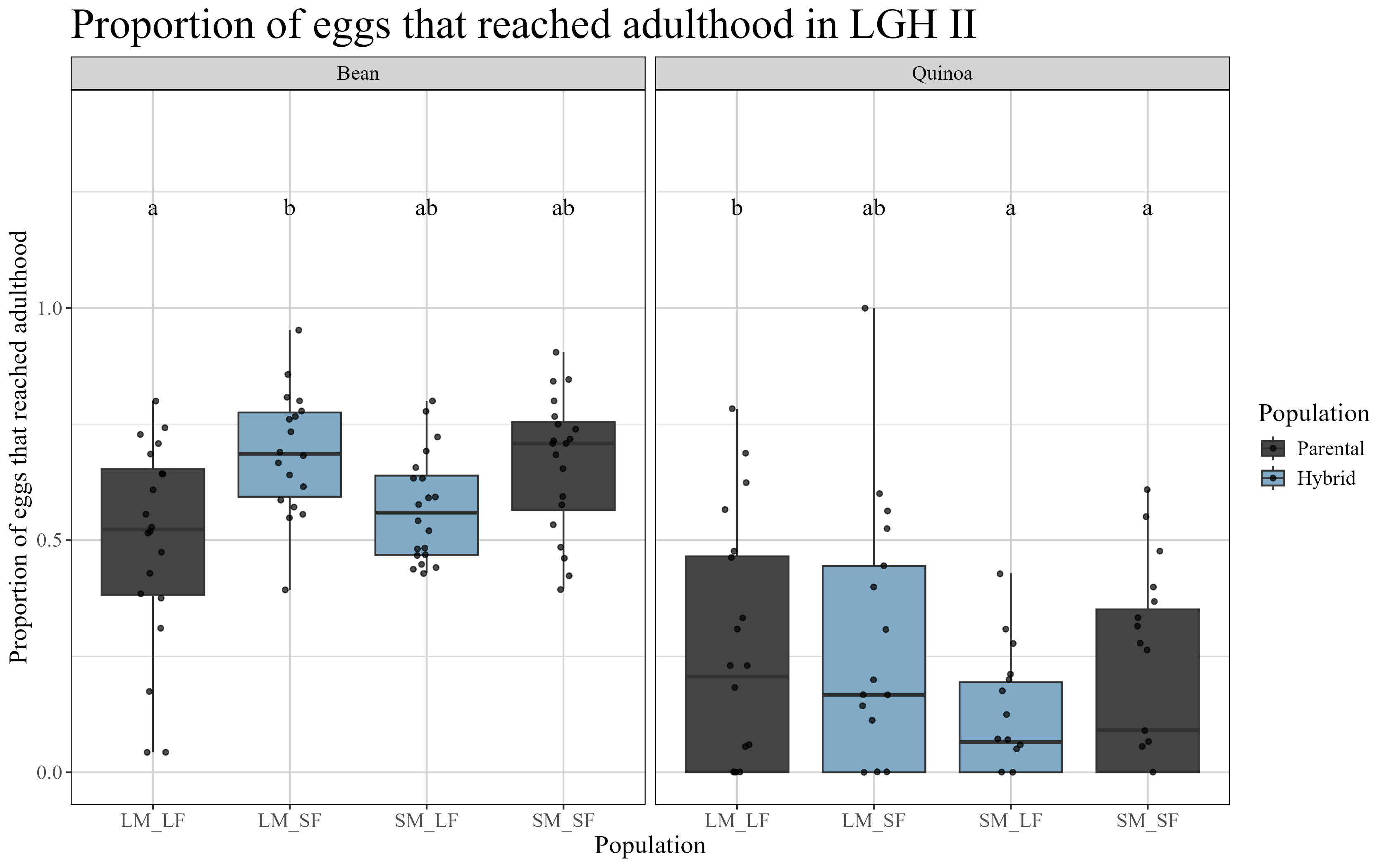

A

B

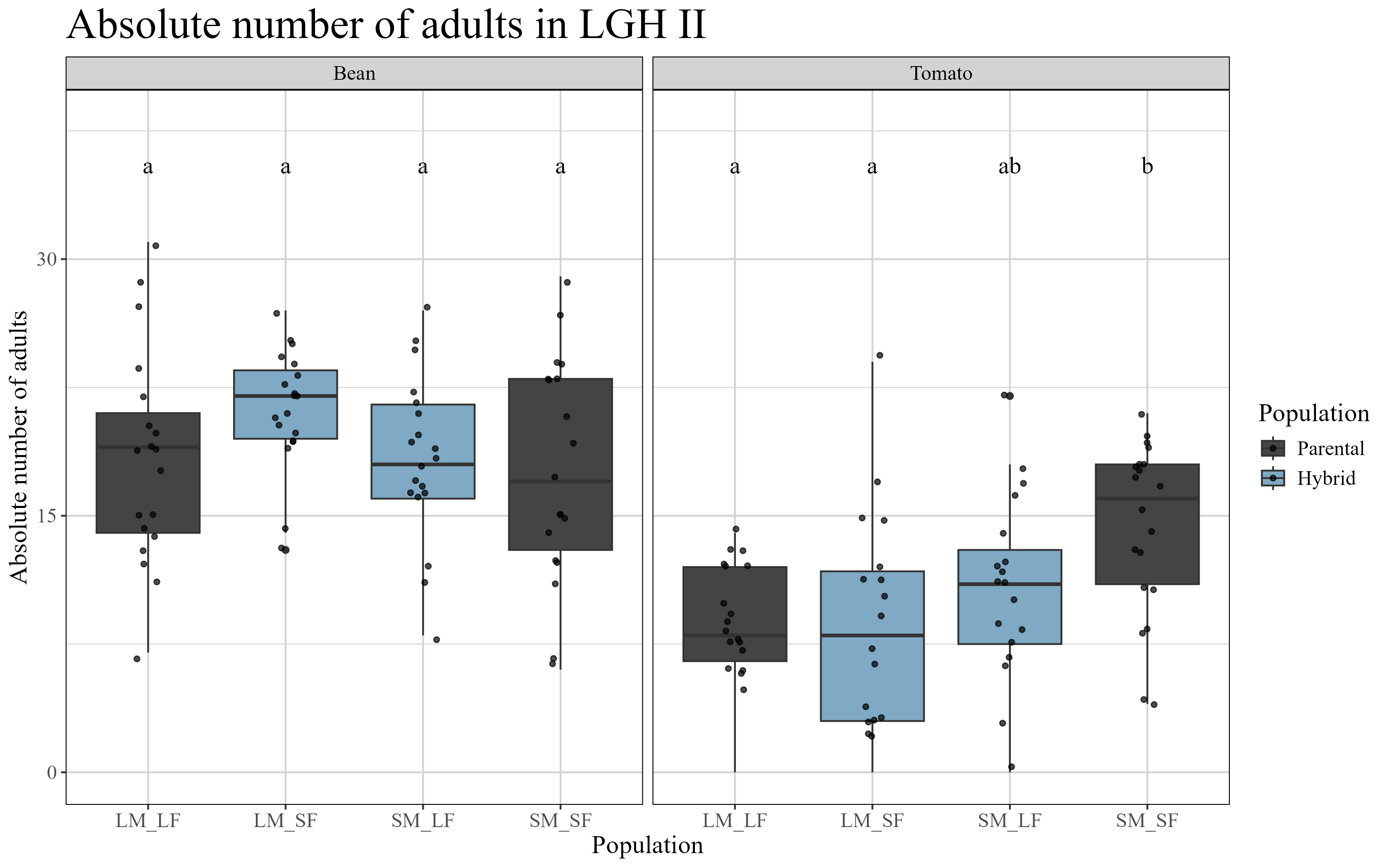

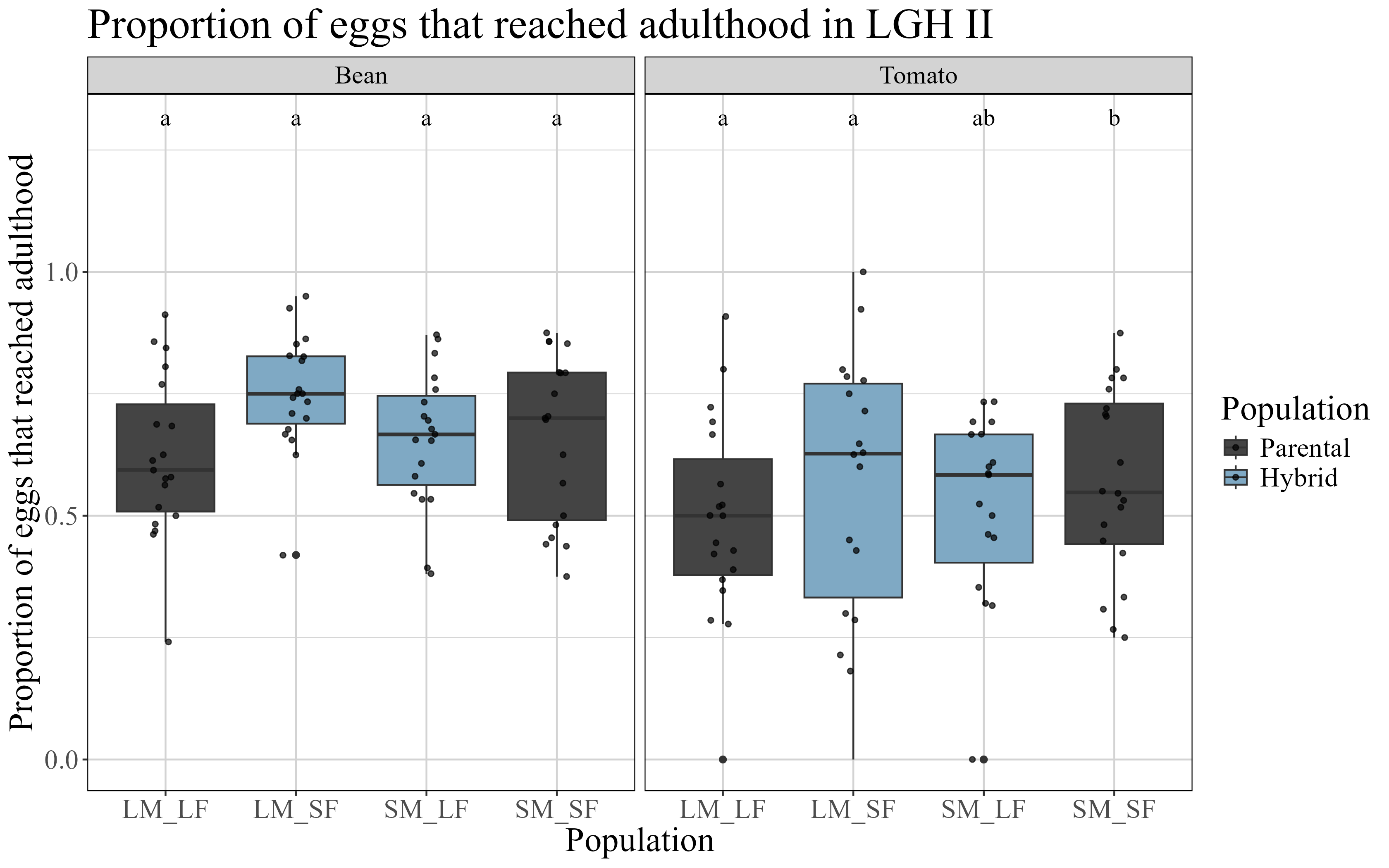

C

D

E

F

Supplementary Figure 3: Fitness measures of parental lines (LMxLF and SMxSF) and Late Generation Hybrids II (LGH II) visualised for both crossing directions (LMxSF and SMxLF) tested on the ancestral (bean) and three different novel host plants: a + b) quinoa, c + d) maize and e + f) tomato. Left panels (a, c, and e) show the absolute number of emerged adults. Right panels (b, d, and f) show the ratio of eggs that reached adulthood, calculated as the number of adults divided by the number of eggs initially laid (n = 20 per treatment). Different letters indicate statistically significant differences (p < 0.05).

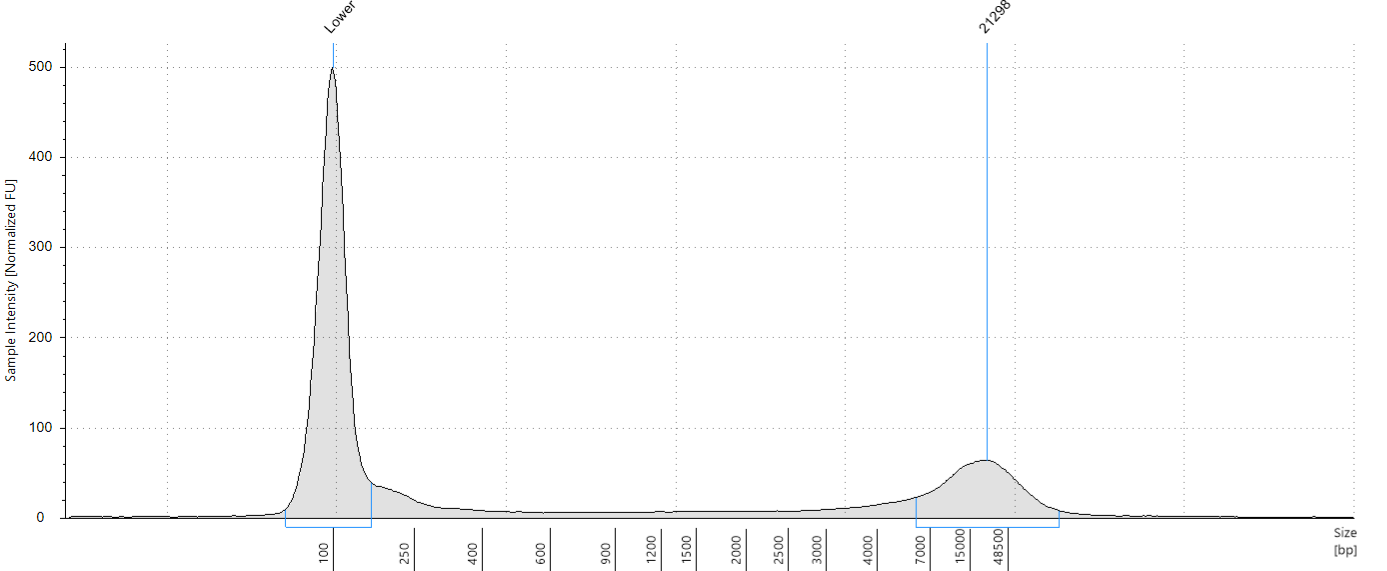

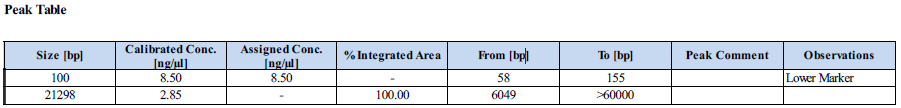
 Supplementary Figure 4. Quality and concentration assessment of single spider mite genomic DNA. Representative electropherogram from an Agilent 4150 TapeStation run using 1 μl of genomic DNA (gDNA) extraction from a single *Tetranychus urticae* female individual.

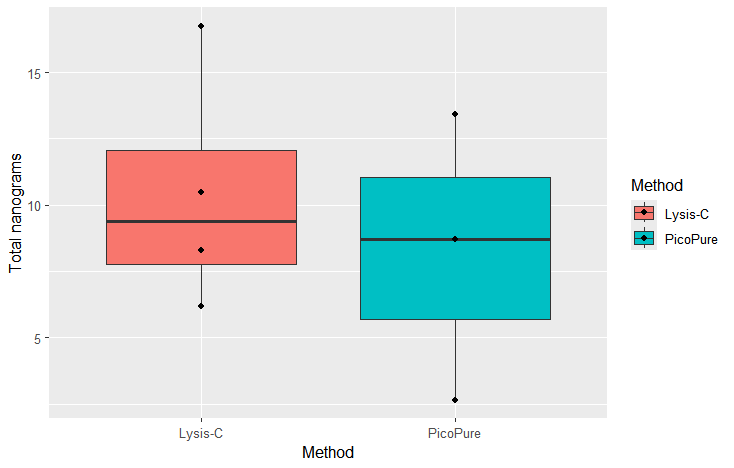

Supplementary Figure 5. Comparison of total genomic DNA obtained from single mites using the 'SOP – Lysis C plate-based DNA extraction' (Korlević et al., 2023) and the PicoPure DNA Isolation Kit (ThermoFisher). No significant difference in DNA yield was detected between the two methods (Two-sample t-test; t = 0.56, df = 4.0, p = 0.6).
